# Pollution-Driven Selection in Riparian Ecosystems: Genome-Wide Responses to *Bacillus thuringiensis israelensis* and Copper in a Non-biting Midge

**DOI:** 10.1101/2025.06.30.662114

**Authors:** Nina Röder, Sara Kolbenschlag, Sebastian Pietz, Reid S. Brennan, Mirco Bundschuh, Markus Pfenninger, Klaus Schwenk

## Abstract

Riparian ecosystems are vital interfaces between aquatic and terrestrial environments but are increasingly impacted by anthropogenic pollution. In these systems, merolimnic insects serve as crucial ecological links, occupying aquatic habitats as larvae and terrestrial environments as adults, thus being an essential food source in both. Consequently, pollutant exposure during the aquatic larval stage can have cascading effects across ecosystem boundaries. While the ecological consequences of such exposure are well documented, the evolutionary potential of merolimnic insects to adapt to chronic pollution remains poorly understood. To address this, we previously conducted a selection experiment exposing populations of the non-biting midge *Chironomus riparius* to the mosquito larvicide *Bacillus thuringiensis israelensis* (Bti) or heavy metal copper over approximately eight generations, which revealed only limited evidence of consistent phenotypic adaptation. Here we use whole-genome sequencing of these populations to assess their genomic responses to chronic pollutant exposure. Despite similar phenotypic sensitivity in pre-exposed and naïve populations, we detected distinct stressor-specific genomic responses. Copper exposure induced a significant genome-wide reduction in nucleotide diversity and evidence of selection-driven allele frequency changes, while Bti effects were dominated by heterogeneous, replicate-specific shifts, potentially reflecting drift or selection on multiple redundant pathways. Functional enrichment analyses indicated early-stage adaptation: immune- and apoptosis-related pathways were enriched under Bti, while metal detoxification and DNA repair pathways were enriched under copper, highlighting distinct adaptive mechanisms despite weak genome-wide signals of selection. Our findings demonstrate that Evolve and Resequencing approaches enable the detection of early genomic signals of adaptation even when phenotypic change is subtle or absent, offering a powerful framework for studying evolutionary responses to environmental pollution.

## Introduction

Riparian ecosystems are ecologically rich and dynamic transition areas between land and water. They support high biodiversity and contribute to key ecosystem functions such as nutrient cycling, habitat provision, and the regulation of water quality (Singh et al., 2021). Many organisms in these systems link aquatic and terrestrial environments both physically – by moving between habitats, transporting energy and nutrients – and functionally, through their ecological roles in both habitats and across their life stages (Polis et al., 1997). A key example is the group of merolimnic insects, which develop as aquatic larvae and emerge as terrestrial adults. However, riparian ecosystems are increasingly affected by anthropogenic pressures (Feld et al., 2018; Lind et al., 2019). In addition to well-recognized stressors such as invasive species and hydromorphological changes, anthropogenic pollutants are now widely acknowledged as significant threats to riparian ecological integrity (Schulz et al., 2024).

Anthropogenic pollutants in riparian ecosystems originate from various human activities, including agriculture, pest control, mining, and urban infrastructure. These pollutants include both chemical compounds, such as heavy metals, and biological agents like microbial pesticides. *Bacillus thuringiensis israelensis* (Bti), a microbial larvicide widely applied to surface waters for mosquito control (Brühl et al., 2020), is intentionally introduced into aquatic ecosystems, raising concerns about its effects on other aquatic invertebrates like non-target insect larvae. In contrast, heavy metal copper enters freshwater systems unintentionally but consistently through surface runoff and soil erosion linked to agricultural and industrial activity (Comber et al., 2023; Pesce et al., 2025). It accumulates in sediments, where it remains bioavailable and toxic (Flemming & Trevors, 1989). This poses a significant risk to benthic organisms such as merolimnic insect larvae, which often inhabit or burrow into sediments.

Among the organisms affected by riparian pollutants are non-biting midges of the family Chironomidae, which are particularly important due to their high abundance and crucial roles in nutrient cycling and food web dynamics (Armitage et al., 1995). Chironomid larvae typically inhabit aquatic sediments in large numbers, where they serve as a major food source for aquatic predators (Serra et al., 2016). As they emerge from the water, chironomids often form large, synchronized swarms, becoming a predictable and energetically valuable resource for a variety of terrestrial predators, including birds, bats, and spiders (Baxter et al., 2005; Martin-Creuzburg et al., 2017). Consequently, disruptions to chironomid populations during their larval stage trigger cascading effects across both aquatic and terrestrial food webs in riparian ecosystems (Allgeier et al., 2019; Fukui et al., 2006; Kato et al., 2003; Kolbenschlag et al., 2023).

Bti-based larvicides are designed and applied to be relatively target-specific (Boisvert & Boisvert, 2000). Bti toxins act by binding to specific receptors in the gut epithelium of aquatic dipteran larvae, causing gut disruption and ultimately larval death (Bravo et al., 2007). Despite its specificity, a considerable number of studies have reported adverse effects on chironomid larvae when exposed to Bti during mosquito control practices (Land et al., 2023). For example, mesocosm experiments have shown reductions in chironomid larvae abundance by up to 87% (Allgeier et al., 2019). Bti-induced chironomid reductions have been reported to propagate through aquatic and terrestrial food webs, affecting predators such as newts, dragonflies, damselflies, and insectivorous birds (Allgeier et al., 2019; Jakob & Poulin, 2016; Poulin et al., 2010). Thus, despite its targeted mode of action, Bti may disrupt riparian ecosystem interactions across aquatic and terrestrial boundaries.

In contrast to the relatively specific biological action of Bti toxins, sediment-bound copper exhibits a broad mode of toxicity, affecting multiple physiological processes across a wide range of benthic organisms (Roman et al., 2007). While copper is essential in trace amounts, elevated concentrations cause oxidative stress and perturb membrane integrity, protein function, and other vital cellular processes (Gaetke & Chow, 2003). In chironomids, copper exposure has been linked to reduced survival, delayed emergence, developmental deformities, and DNA damage (Bernabò et al., 2017; Marinković et al., 2011; Martinez et al., 2003). On the population level, copper has also been shown to impair reproductive timing, potentially affecting mating success (Servia et al., 2006). These findings underscore that sediment-associated copper can undermine chironomid life-history traits from molecular to population scales, posing a serious threat to benthic community stability.

While the ecological effects of Bti and copper on non-target chironomids are well documented, much less is known about the insects’ capacity for adaptation to these pollutants. Resistance to individual Bti toxins has been demonstrated in some mosquito populations (Paris et al., 2011a; Tetreau et al., 2013), but there is currently no evidence for adaptation to commercial Bti formulations in either mosquitoes or chironomids (Becker et al., 2018; Brühl et al., 2020; Tetreau et al., 2013). In contrast, *C. riparius* has shown some potential for evolutionary responses to metal exposure. Early studies documented reduced sensitivity in historically contaminated (Groenendijk et al., 1999a; Postma et al., 1995a; Postma et al., 1995b), cross-bred adapted, (Groenendijk et al., 2002) or laboratory-selected lines (Marinković et al., 2012; Postma & Davids, 1995; Vogt et al., 2007), though these often lacked controls for maternal effects, phenotypic plasticity and/or unintended adaptation to laboratory conditions, making it difficult to attribute observed phenotypic changes strictly to genetic adaptation (Doria et al., 2022; Marinković et al., 2012). More recently, Im et al. (2019) provided physiological and epigenetic evidence of heritable adaptation to metal-contaminated sediments, and Doria et al. (2022) detected clear genomic signatures of cadmium tolerance in *C. riparius* after eight generations of laboratory exposure. However, robust evidence for copper-specific adaptation in *C. riparius* is currently lacking. Understanding the potential for genetic adaptation is essential for predicting population resilience and long-term ecological consequences of anthropogenic pollution in riparian ecosystems.

To shed light on this issue, we previously conducted a selection experiment with the non-biting midge *Chironomus riparius,* an established model species in ecotoxicology and evolutionary ecology (Foucault et al., 2019; Liu et al., 2025). We exposed populations to Bti or copper for approximately eight generations (Kolbenschlag et al., 2024; Pietz et al., submitted). Although some pre-exposure effects were observed – such as altered emergence timing or lipid content at certain Bti concentrations, and increased emergence success at one of the tested copper concentrations – evidence for consistent phenotypic adaptation was limited (Kolbenschlag et al., 2024; Pietz et al., submitted). Here, we build on this experiment by assessing genomic responses to chronic pollutant exposure in the same populations. Investigating allele frequency changes that may underlie adaptive processes allows us to explore potential genomic changes that may not have yet been reflected in the phenotype, for example due to the relatively short duration of the chronic exposure (cf. Doria et al., 2022). Specifically, this study addresses three key objectives: First, we aim to characterize genome-wide changes in *C. riparius* populations exposed to two distinct anthropogenic pollutants — a biologically targeted microbial pesticide vs. a broadly acting metal pollutant — and to compare these with untreated control populations. Second, we seek to determine the specific contribution of selective pressures imposed by these stressors to the observed genomic changes, distinguishing them from neutral processes such as genetic drift. Third, we investigate the functional basis of these genomic changes by identifying biological processes significantly overrepresented among affected genes, thereby uncovering the evolutionary responses triggered by each stressor. By linking pollutant-specific selective pressures to genomic signatures and associated biological functions, this study provides insights into how anthropogenic pollution may drive evolutionary change in riparian ecosystems.

## Material and methods

### Laboratory population establishment and culture conditions

To increase genetic diversity in the study population before chronic exposure, *C. riparius* in-house cultures were mixed with individuals of the same species from the laboratories of BASF SE (Ludwigshafen, Germany), ECT Oekotoxikologie GmbH (Flörsheim, Germany) and the LOEWE Centre for Translational Biodiversity Genomics (Frankfurt am Main, Germany) in the four months prior to the experiment. This approach was based on the assumption that long-established laboratory cultures may lack sufficient genetic variation to exhibit responses to selective pressures (Nowak et al., 2012). To minimize variability during the experiment due to newly introduced genotypes, no further individuals were added during the two months preceding the start of the exposure, allowing the laboratory population to stabilize. All cultures had been maintained for several months to years prior to their use in this experiment, following standard protocols for lab-based toxicity testing (e.g., OECD guidelines). Accordingly, food and sediment were provided for the aquatic larvae in water containers, which were placed inside mesh cages tall enough to allow adult emergence and successful reproduction. Climate chambers were used to maintain stable temperature, humidity, and controlled day-night cycles. Although a new generation can appear in as little as two weeks (OECD, 2004), the exact number of generations that occurred within a longer time period is uncertain, as generation times can vary, with overlapping generations being likely (Foucault et al., 2019).

The experimental setup was conducted within three identical climate chambers, maintaining a temperature of 20±1 °C, 65% humidity, and a 16:8 day/night rhythm. Each experimental unit comprised a cage (50 x 35 x 50 cm, L x H x W, mesh size: 0.6 mm) housing two test vessels (32 x 7 x 22 cm). Within each vessel, there was 1.1 kg (wet weight) standardized sediment (composed of 75% sand, 20% clay, 5% peat; 40% water and 0.1% CaCO_3_), and 2 L gently aerated SAM-5S medium (Borgmann, 1996), in accordance with the respective OECD guidelines for chironomid toxicity testing (OECD, 2010, 2023).

### Egg collection, larval distribution, and chronic exposure

Between April 21 and 28, 2021, a total of 84 egg masses, each consisting of several hundred eggs, were collected from the culture over four days and stored in SAM-5S medium. Typically, *C. riparius* larvae begin to hatch 2 to 3 days after the eggs were laid at 20 °C (OECD, 2004). On days 4 and 6 after the end of the collection period, the freshly hatched larvae were split into 21 groups, with the aim of achieving an even number of individuals per replicate. Because each egg mass contains several hundred individuals, it is not feasible to individually count and allocate individuals manually to replicates, as the handling and the duration of the process might harm the larvae. Instead, we used a custom-built rotary dispensing device designed to ensure uniform distribution of larvae. This device consisted of a rotating basket with 21 glass tubes evenly distributed along the outer edge, paired with a pipette equipped with a funnel. By pouring the larval medium in a thin stream while rotating the basket, larvae were evenly dispensed into each tube. To estimate the numbers of larvae in each tube, three of the samples were individually counted under a stereomicroscope using pipettes modified with glass-tips. On day 4, these counts yielded an average of 350 larvae per replicate (range: 347–358), and on day 6, approximately 50 larvae per replicate (range: 45–60), resulting in a total of approximately 400 larvae per replicate. The remaining 18 replicates were assumed to contain similar numbers and were each allocated to one of 18 experimental units. Each group (day 4 and day 6) was introduced into one of the two vessels prepared for each experimental unit. The larvae used for counting were preserved in 70% ethanol at −20 °C to represent the genomic structure of the founding population, serving as a baseline for comparison with individuals sampled after chronic exposure.

The experimental conditions consisted of six replicates each of a control, a Bti treatment and a copper treatment. Treatment concentrations were selected based on preliminary tests (data not shown), with the goal of applying sufficient selection pressure without causing populations to collapse. In the Bti treatment, each experimental unit was treated every two weeks with Bti using the commercial formulation VectoBac WDG (Valent BioSciences, Illinois, USA) at 33% of the recommended field rate, corresponding to 480 ITU/L. Repeated application was necessary to ensure comparable exposure of each chironomid generation, as the relatively high water temperatures, water turbidity and the absence of vegetation likely limited the duration of Bti toxicity (Brühl et al., 2020). Furthermore, it was assumed that the largest fraction of Bti toxins was lost during the biweekly test medium exchange (see below). The copper treatment involved sediment spiked with copper sulfate to achieve an environmentally relevant nominal concentration of 100 mg Cu/kg dry weight. These sediments were prepared and covered with medium two weeks prior to larval introduction to allow copper concentrations to equilibrate between sediment and porewater (OECD, 2004; Simpson et al., 2004). No additional copper was added during the chronic exposure phase, under the assumption that the majority remained bound to the sediment. The remaining six cultures served as untreated controls. To maintain stable water quality, the test medium was renewed every second week, within 48 hours prior to Bti application. Evaporation was compensated for by regularly refilling the vessels to maintain a constant water level. Larvae were fed ground TetraMin fish food (Tetra GmbH, Melle, Germany) twice a week at a rate of approximately 0.5 mg per larva per day.

After 26 weeks, about six months, of chronic exposure to copper and Bti (13 Bti applications), egg masses from each experimental unit were sampled on three consecutive days and stored in SAM-5S medium until hatching, resulting in a median of 10 egg masses per replicate (range: 4–22; SD = 5.6; see Supporting Information, Figure S1). Three days later, 100 larvae from each experimental unit were counted and stored in 70% ethanol at −20 °C for subsequent genomic analysis. Although generations likely overlapped during the chronic exposure period, we estimated that the sampled larvae represent approximately the eighth generation, based on an average generation time of three to four weeks.

### DNA extraction and whole-genome sequencing

We used 100 larvae of each experimental unit (n = 18) and 400 larvae from the founding population, the latter of which was split into two technical replicates during DNA extraction. Following careful removal of ethanol from the samples, larvae were dried at 56 °C for approximately 4 hours. DNA extraction was carried out using the QIAamp DNA Mini Kit (Qiagen, Hilden, Germany) following the manufacturer’s instructions, involving a 2-hour proteinase K lysis step and elution in 25 µL elution buffer. Library preparation with a 450 bp insert size and subsequent sequencing of 150 bp paired-end reads were performed by Novogene Europe – UK on an Illumina NovaSeq platform, aiming for an expected coverage of 30x.

### Single nucleotide polymorphism (SNP) identification

Reads were trimmed for quality and adapter contamination using trimmomatic v. 0.39 (Bolger et al., 2014) and then mapped on the latest *C. riparius* reference genome on NCBI (Bioproject number PRJEB47883) using BWA mem (Li & Durbin, 2009). Before trimming, the two technical replicates of the founding population were merged into one file. Variants were called using VarScan 2 (Koboldt et al., 2012) with a minimum variant frequency of 0.01, P value of 0.1, minimum alternate reads 2, and minimum coverage of 30x, resulting in 8,498,585 sites. Sites were then filtered for only biallelic sites with a coverage of >30x in all samples and a minimum minor allele frequency of 0.05 in at least six samples (i.e., one treatment). To control for mismapping, sites with depth per sample above the 97.5% quantile (951x for the founding population and 416x for all other populations) were excluded. The filtering resulted in a final set of 119,556 biallelic SNP sites.

### Characterizing genomic variation

To assess the population genomic effects of the selection regimes, we estimated genome-wide levels of nucleotide diversity and linkage disequilibrium (LD) for all replicates. Nucleotide diversity was estimated as Tajima’s pi (π) in the founding population and each evolved replicate using PoPoolation (Kofler et al., 2011a). We estimated π in 100-bp sliding windows with a 100-bp step size, resulting in 1,918,381 100-bp windows in 4 chromosomes and 10 scaffolds across the genome that were present across all samples. Each window required a minimum coverage of 25x, maximum coverage of 1,000x (to avoid mapping errors), and at least half of the window meeting these thresholds. To test for differences in π among treatments while accounting for replicate structure, we used a linear mixed-effects model (LMM) with treatment as a fixed effect and replicate as a random effect. The model was fitted using the *`lmer`* function from the `lme4` R package (Bates et al., 2015). To assess treatment differences, we performed pairwise comparisons of estimated marginal means using the `emmeans` package (Lenth, 2025), applying Holm correction for multiple testing. All statistics were performed in R (R Core Team, 2024). We summarized genome-wide variation in allele frequencies between all samples using principal component analysis (PCA) with the *`prcomp`* function, following an arcsine square root transformation of the allele frequency data to stabilize variances. LD was estimated using LDx, a Pool-Seq method that uses haplotype information from single read pairs to estimate linkage between pairs of SNPs over short distances (Feder et al., 2012). We estimate the decay of LD by regression of the log of physical distance with LD between base pairs; to estimate the slope and intercept of each treatment, we include replicate as a random effect with the R package `nlme` (Pinheiro et al., 2024).

### Estimates of allele frequency change within and between treatments

We determined specific loci evolving due to selection by simulating the expected drift over eight generations using the `Pool-Seq` package in R (Taus et al., 2017) and following Barghi et al. (2020). Using the starting allele frequencies at F0 (the founding generation), we mirrored our experimental design and simulated allele frequency trajectories for six replicates across eight generations with no selection under a Wright-Fisher model. We used the `Pool-Seq` R package to estimate the mean effective population size as 154 across all replicates, and simulated neutral allele frequency changes using this effective population size and a census size of 600. The census size is defined as the average number of reproducing individuals. Although individual counts were not tracked precisely, we estimated that each cage replicate supported approximately 600 reproducing individuals per generation based on observed densities and the assumption that each chironomid typically requires about 2 cm² of sediment surface to develop (OECD, 2004). We added the same variance as our sampling scheme by estimating allele frequencies from a sample size of 100 individuals and a simulated sequencing depth of 136x at F0 and 68x at F8 (the final generation), matching the observed depth. This simulation was repeated 500 times and, for each replicate, Fisher’s exact tests in PoPoolation2 (Kofler et al., 2011b) were used to generate a null distribution of neutral allele frequency change. Empirical P values at the level of individual loci were calculated from this simulated distribution using *`empPvals`* in the `qvalue` R package (Storey et al., 2024), which estimate the proportion of Fisher’s exact test statistics from the neutral simulations that are equal to, or more extreme than, those observed in the experimental data. This approach identifies loci where the observed allele frequency change is unlikely under genetic drift alone, thereby highlighting candidates for selection. Given that polygenic adaptation often involves genetic redundancy, where different replicate populations can reach similar phenotypic outcomes via distinct genetic routes (cf. Barghi et al., 2019), we did not require all replicates to show identical signals of selection. Candidate adaptive loci were sites with empirical P values less than 0.05 in at least four out of six replicates per treatment i.e., at least four replicates showed SNP variation exceeding expectations under genetic drift. Significance of overlap between sets of candidate SNPs between treatments was calculated using SuperExactTest in R (Wang et al., 2022).

### Disentangling selection, drift, and laboratory adaptation in allele frequency changes

When selection acts on standing genetic variation in polygenic traits, the resulting allele frequency changes are often subtle and difficult to distinguish from genetic drift (Berg & Coop, 2014). To improve the detection of such signals, we applied a covariance-based approach that accounts for confounding effects like drift and laboratory adaptation (Buffalo & Coop, 2020), allowing us to quantify the contribution of selection to the variance in allele frequency change within and between treatments (cf. Brennan et al. 2022). This method partitions the variance in allele frequency change from F0 to F8 into components attributable to selection, drift, and lab adaptation, based on covariance across replicate populations within treatments. Covariance in allele frequency change was computed in 10,000-bp windows along the genome for each replicate. Uncertainty was assessed through bootstrap resampling of windows. Temporal changes in allele frequencies that were consistent across at least four out of six replicates within a treatment were defined as the shared response for that treatment. To estimate the portion of this shared response attributable to selection, we subtracted the component attributable to laboratory adaptation. Since the control is expected to be selected only for culturing conditions, any covariance observed between the control and the treatment populations was interpreted as adaptation to the laboratory environment. When calculating shared responses based on subsets of four or five replicates, we conservatively identified the corresponding subset of control replicates (i.e., four or five out of six) that yielded the highest covariance with the treatment subset. We used this maximum covariance to quantify lab adaptation. Treatment-specific parallel changes were only attributed to selection if they exceeded this lab-adaptation signal.

### Functional enrichment analysis

We used InterProScan (Jones et al., 2014) to assign Gene Ontology (GO) terms to predicted protein sequences. GO enrichment analysis was performed using topGO (Alexa & Rahnenfuhrer, 2024), applying the weight01 algorithm to identify significantly enriched biological processes. GO terms with fewer than five annotated genes were excluded by default during analysis. Prior to enrichment analysis, we filtered SNPs based on significance (false discovery rate < 0.05) in at least four, five, or all six replicates for each treatment. Then, topGO was run separately for each treatment group to assess treatment-specific GO term enrichment. A GO term was considered significant if at least one SNP associated with that GO term passed the threshold (Fisher statistic < 0.05). GO terms hierarchical information was retrieved from AmiGO2 (Carbon et al., 2009) and biological processes unique to treatments were identified. REVIGO (Supek et al., 2011) was used to generate a two-dimensional visualization of GO terms based on their semantic similarity, reflecting the degree of relatedness in their biological annotations.

## Results

### Genome-wide variation in response to experimental selection

To assess genome-wide responses to anthropogenic pollutants, we compared allele frequency changes in *C. riparius* populations exposed to chronic Bti or copper treatment, with unexposed controls and the founding population. We used pool-sequencing data from each replicate, yielding 119,556 SNPs with no missing data across all samples. The founding population possessed genome-wide genetic diversity on which selection could act (Tajima’s π: 0.008 ± 0.010). Principal-component analysis (PCA; Figure 1) showed that the variance in genome-wide allele frequencies for all samples did not obviously cluster by treatment group, and not all F8 samples had notably diverged from the F0 founding population along PC1 (12.3% of the variation). There was no clear separation of the treatments from the control along this axis. However, four replicates — two from copper-exposed populations and one each from the Bti-treated and control populations — were positioned slightly further apart from the rest along PC1. Along PC2 (8.4% of the variation), two of these replicates (one Bti- and one copper-treated population) also appeared further apart from the remaining samples, although this axis likewise did not group treatments into distinct clusters.

**Figure 1.**
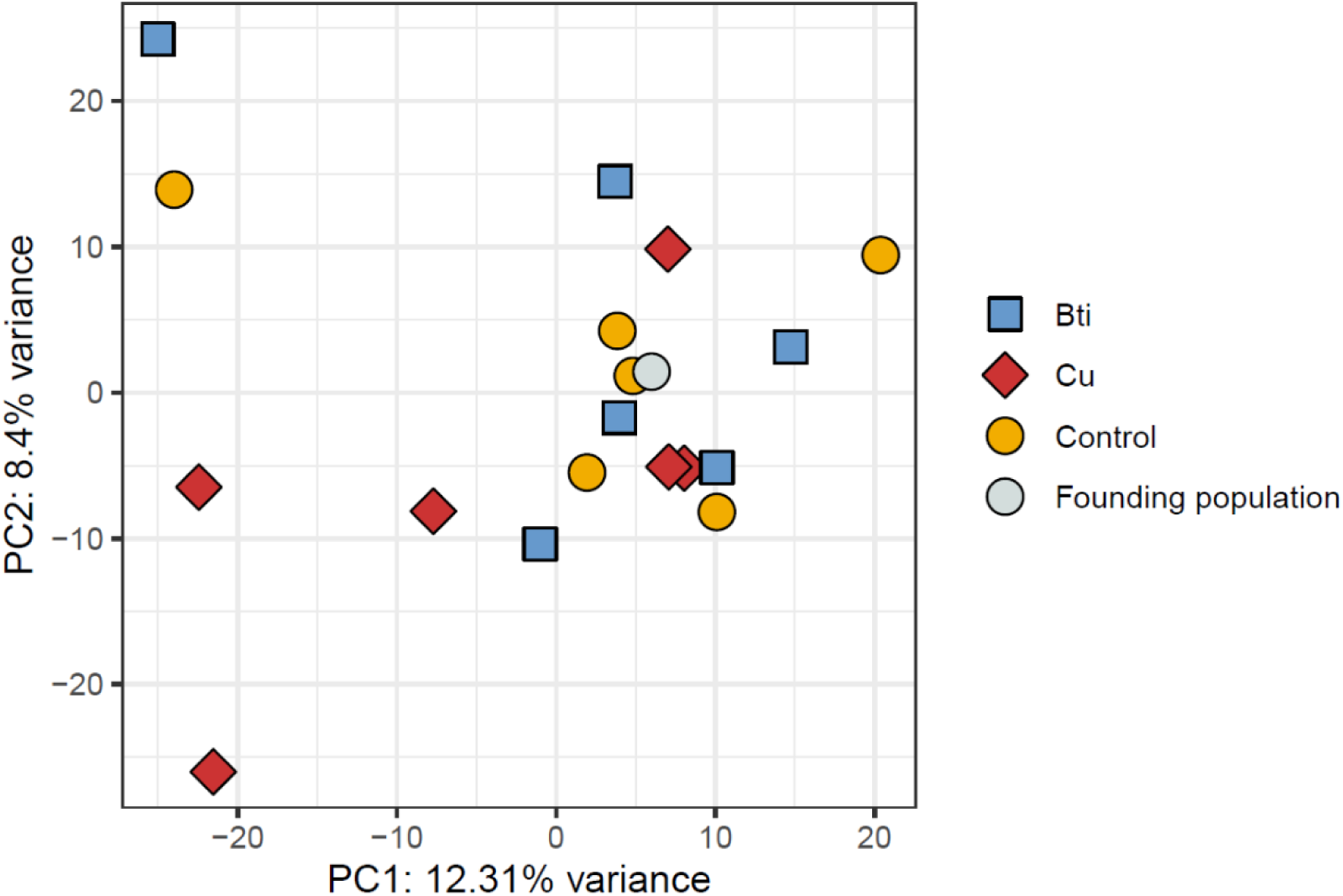
PCA of allele frequencies from 119,556 SNPs across the genome where color and shape distinguish treatment groups. The founding population is F0; all others are from F8.

We observed a slower rate of LD decay in the F8 populations compared to the founding population, indicating that all F8 populations underwent a bottleneck (Founding population: intercept: 0.3047 ± 0.0006; slope: −0.0447 ± 0.0003). The copper-exposed populations showed the largest increase in LD (intercept: 0.3096 ± 0.0036; slope: −0.0361 ± 0.0001), followed by Bti (intercept: 0.3101 ± 0.0039; slope: −0.0390 ± 0.0001) and control (intercept: 0.3113 ± 0.0022; slope: −0.0404 ± 0.0001). Because LDx estimates are sensitive to parameter settings (Feder et al., 2012), the LD values should be interpreted as relative differences between the treatments rather than exact measures of LD.

While adaptation to environmental stressors can promote population persistence, selection due to pollutant exposure typically reduces standing genetic variation. Consistent with this expectation, genome-wide nucleotide diversity (Tajima’s π) decreased by between 4.8% and 8.4% across all treatment groups relative to the founding population. The greatest reduction was observed in the copper treatment (8.4% decrease; π = 0.0076 ± 0.0010; LMM p = 0.04), followed by the Bti treatment (6.0% decrease; π = 0.0078 ± 0.0011; p = 0.20), and the control populations (4.8% decrease; π = 0.0079 ± 0.0011; p = 0.20).

### Identifying potential selection targets

To pinpoint the genetic variation that may underlie adaptation to the different experimental selection regimes, we simulated the expected level of genetic drift over eight generations based on our experimental design (see *Material and methods*). We then employed Fisher’s exact tests to identify loci that were evolving at a rate significantly higher than what would be expected due to drift alone in each replicate. Candidate adaptive loci were sites with empirical P values less than 0.05 in at least four out of six replicates per treatment i.e., at least four replicates showed SNP variation exceeding expectations under genetic drift. We found 1,932 (1.6% of all SNPs), 1,551 (1.3%), and 1,706 (1.4%) SNPs that were candidate targets of selection for Bti treated, copper-exposed, and control conditions, respectively (Figure 2). While most loci were unique within treatments, some were shared between treatments. All treatments showed a similarly high level of unique responses, with 1,718 SNPs (88.9% of significant loci) in the Bti-treated populations, 1,357 SNPs (87.5%) in the copper-exposed populations, and 1,495 SNPs (87.6%) in the control populations showing treatment-specific signals (Figure 2). However, the pairwise shared response between treatment groups was greater than by random chance (exact test, P < 0.0001) and similarly high (111 SNPs, 94 SNPs and 91 SNPs) for all pairs of treatments. There was a very small shared response between all treatments (9 SNPs), suggesting a weak signal of shared adaptation to laboratory conditions.

**Figure 2.**
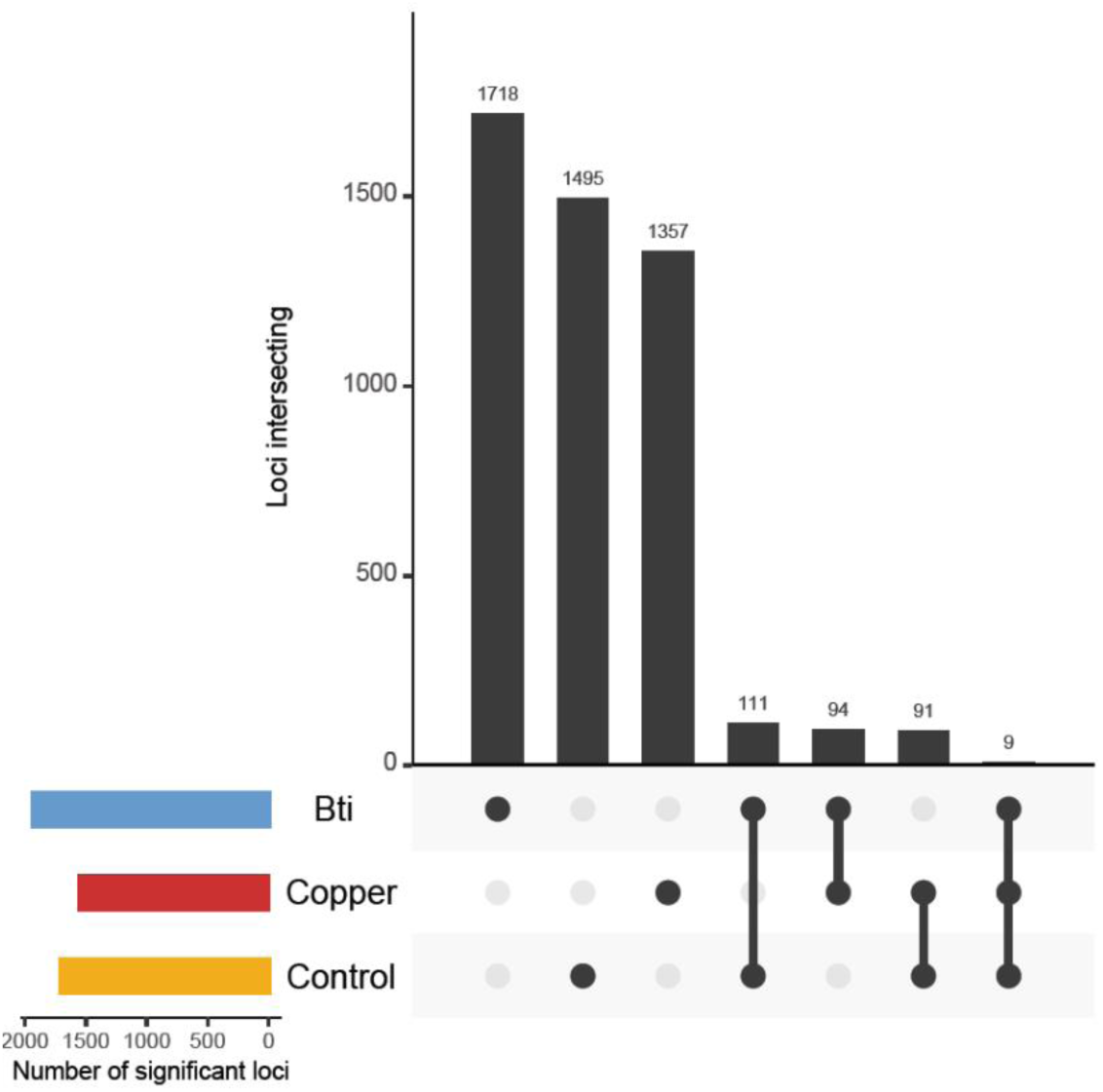
Candidate SNPs exceeding drift expectations identified from Wright–Fisher simulations in at least four out of six replicates. The horizontal bars indicate the total number of candidate loci in each group while the vertical bars show the number in each category. Black points show treatments in a category where multiple groups are indicated by connected lines. Note that the counts in each group are exclusive; for example, the number of loci in the control-alone category are those loci not shared with any other group.

When stricter conditions were applied, i.e., requiring 5 out of 6 replicates per treatment to show significant SNPs, the number of treatment-specific SNPs decreased by a factor of approximately 7, with around 220 SNPs identified for each treatment instead of the ∼1,500 SNPs under the initial condition (Supporting Information, Figure S2). The number of SNPs shared between two treatments also decreased to a range of 1 to 7 SNPs, compared to around 100 under the more lenient condition. Notably, no SNPs were shared across all three treatments. When even stricter conditions were applied, requiring all six replicates to show significant SNPs, the number of candidate SNPs further decreased to 20 for Bti-treated, 7 for copper-exposed, and 11 for non-treatment conditions, with no overlapping SNPs identified across any treatments (Supporting Information, Figure S2).

### Contribution to allele frequency change: selection, laboratory adaptation, drift and replicate-specific responses

By taking advantage of both the replicated and temporal aspects of our experimental design, we applied a temporal covariance-based method to detect even subtle signals of polygenic selection in genome-wide allele frequency changes. Unlike methods such as Cochran-Mantel-Haenszel (CMH), which focus on identifying strong, consistent selection at a limited number of loci, temporal covariance-based approaches can identify more subtle and weak responses to selection (Buffalo & Coop, 2020). In this way, we can assess the extent to which allele frequency changes across replicates are driven by shared selection pressures rather than random genetic drift or replicate-specific variation (Brennan et al., 2022). We calculated pairwise covariances in allele frequency changes from F0 to F8 within 10,000-bp windows across all replicate pairs. Using this, we derived the convergent correlation, a standardized measure of the similarity in allele frequency changes between replicates within or across treatments, reflecting a convergent selection response. A high convergent correlation indicates parallel allele frequency shifts driven by a shared selective pressure. In contrast, if changes were driven by genetic drift or replicate-specific selection, allele frequency shifts would be independent across replicates, resulting in a convergent correlation of zero. Additionally, if selection acts in divergent directions across populations, it generates negative convergent correlations. We observed comparably high convergent correlations across all pairwise replicate comparisons, regardless of whether they were within the same selection regime or across different treatments (Figure 3).

**Figure 3.**
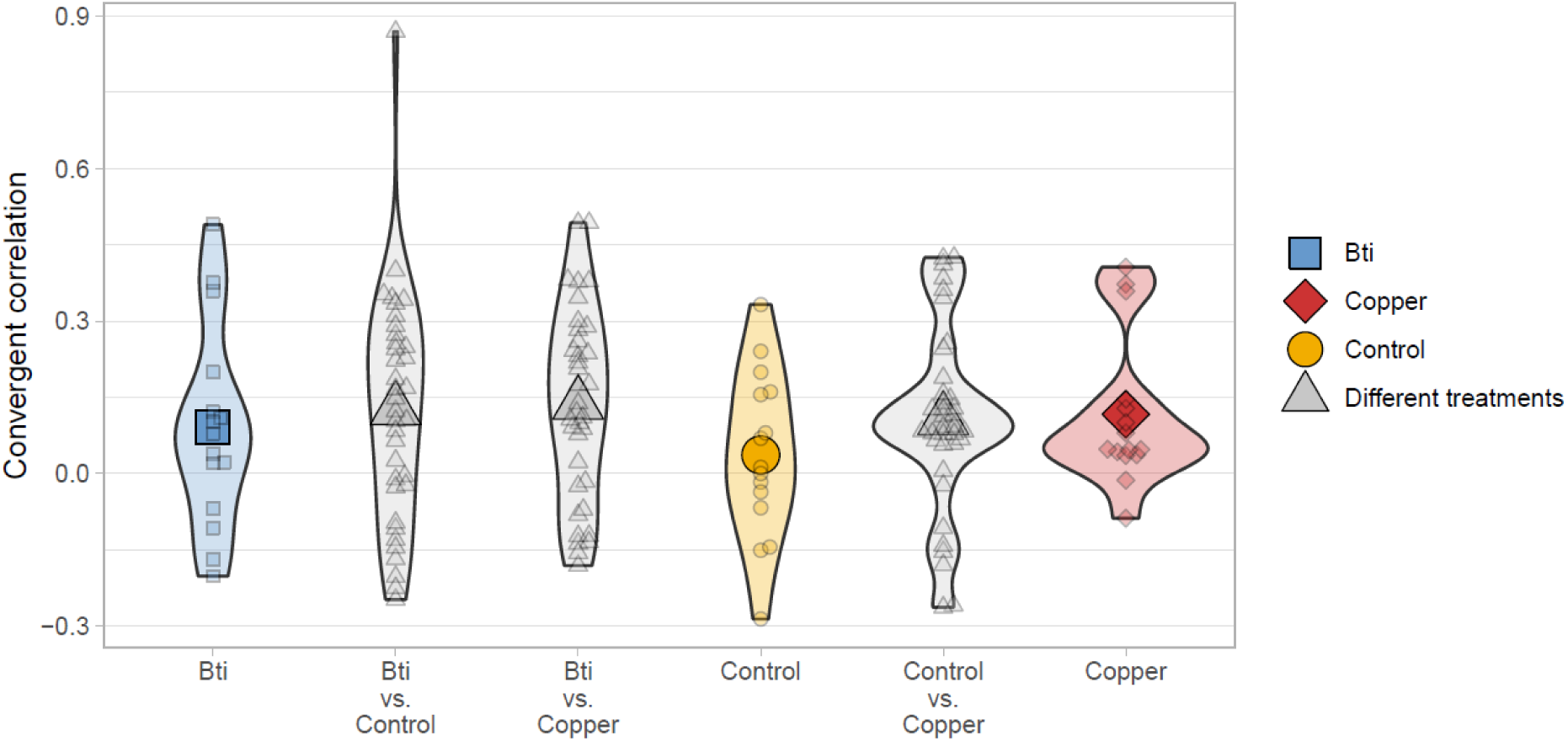
Convergent correlations of allele frequency change from F0 to F8. Higher values indicate a more similar change in allele frequency between two replicates. Small symbols represent convergent correlations from pairwise population comparisons; large symbols show the mean for each group comparison.

We partitioned the total variance in allele frequency change into components attributable to experimental selection, genetic drift, and laboratory adaptation (Figure 4). To conservatively identify signals of selection, we focused on sets of replicates within each treatment that showed a shared response with a positive lower confidence bound — indicating a statistically significant signal above zero. In both the Bti-and copper-treated populations, the strongest shared response was found in specific four-replicate subsets. Within these, the estimated contribution of laboratory adaptation to total variance was higher and more variable in Bti-treated (27%, 95% CI: [5-49%]) than in the copper-exposed populations (19%, 95% CI: [15-22%]). After accounting for lab adaptation, experimental selection explained 0% of the variance in Bti-treated populations (95% CI: [−44-20%]) and 6% in copper-exposed populations (95% CI: [2-11%]). In both treatments, most of the variance was due to drift or replicate-specific responses, with 85% of the total variance in Bti-treated (95% CI: [31-139%]) and 75% in copper-exposed populations (95% CI: [67-83%]; Figure 4).

**Figure 4.**
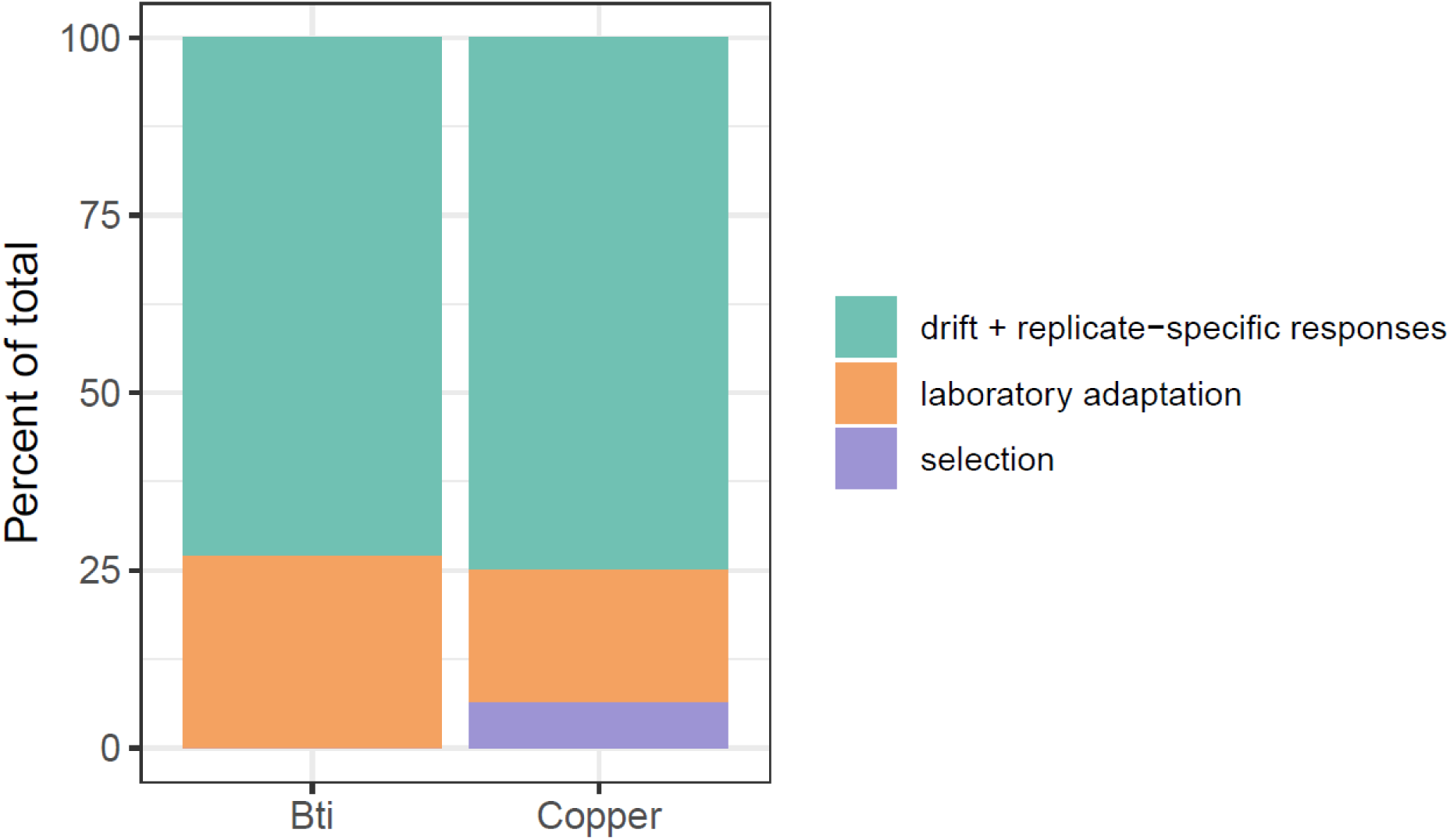
The contribution of laboratory adaptation, treatment selection, and drift plus replicate-specific responses to the total variance of allele frequency change from F0 to F8. The laboratory component was determined using the covariance of allele frequency change between the control and each of the two treatment groups while selection was identified as the covariance within a treatment group minus the laboratory adaptation. The remaining variance was attributed to drift and replicate-specific response to selection.

Across all tested replicate groupings, only four produced statistically significant selection estimates (i.e., lower CI > 0): three four-replicate sets and the full six-replicate set from the copper-exposed populations. Among these, one four-replicate subset slightly outperformed the others, with a selection estimate of 7% (95% CI: [3-10%]). In contrast, no subset from the Bti treatment showed evidence of selection above zero, though the confidence intervals were notably wider than for the copper treatment, reflecting higher variability across genomic windows. Full results for all tested groupings, including estimates of shared response and selection signal, are provided in Supporting Information, Figure S3 and Table S1.

### Gene ontology enrichment

Tests for gene ontology (GO) functional enrichment were used to gain insight into the mechanisms that may underlie adaptation to the different conditions. The GO enrichment analysis identified 45 biological processes associated with loci exhibiting significant allele frequency changes after chronic exposure (Supporting Information, Table S2). Among the treatments, we found enrichment for 11 GO terms in Bti-treated populations, 12 in copper-exposed populations, and 10 in control populations. Additionally, five GO terms were shared between Bti-treated and control populations, three between copper-exposed and control populations, and four were enriched across all treatments. All 45 enriched terms were predominantly related to biological regulation and cellular processes, including metabolic processes and organization or biogenesis of cellular components.

Enrichment of GO terms related to cell structure was primarily observed in control and/or Bti treatments, while those associated with cellular processes were more prominently enriched in the copper and Bti-treated populations. This trend was also evident in the enrichment of GO terms related to stress response, with no specific terms showing enrichment in the control groups. The number of GO terms associated with metabolism was similar across all three treatments, although a distinct separation of clusters by treatment can be observed in the semantic space, as shown in Figure 5.

**Figure 5.**
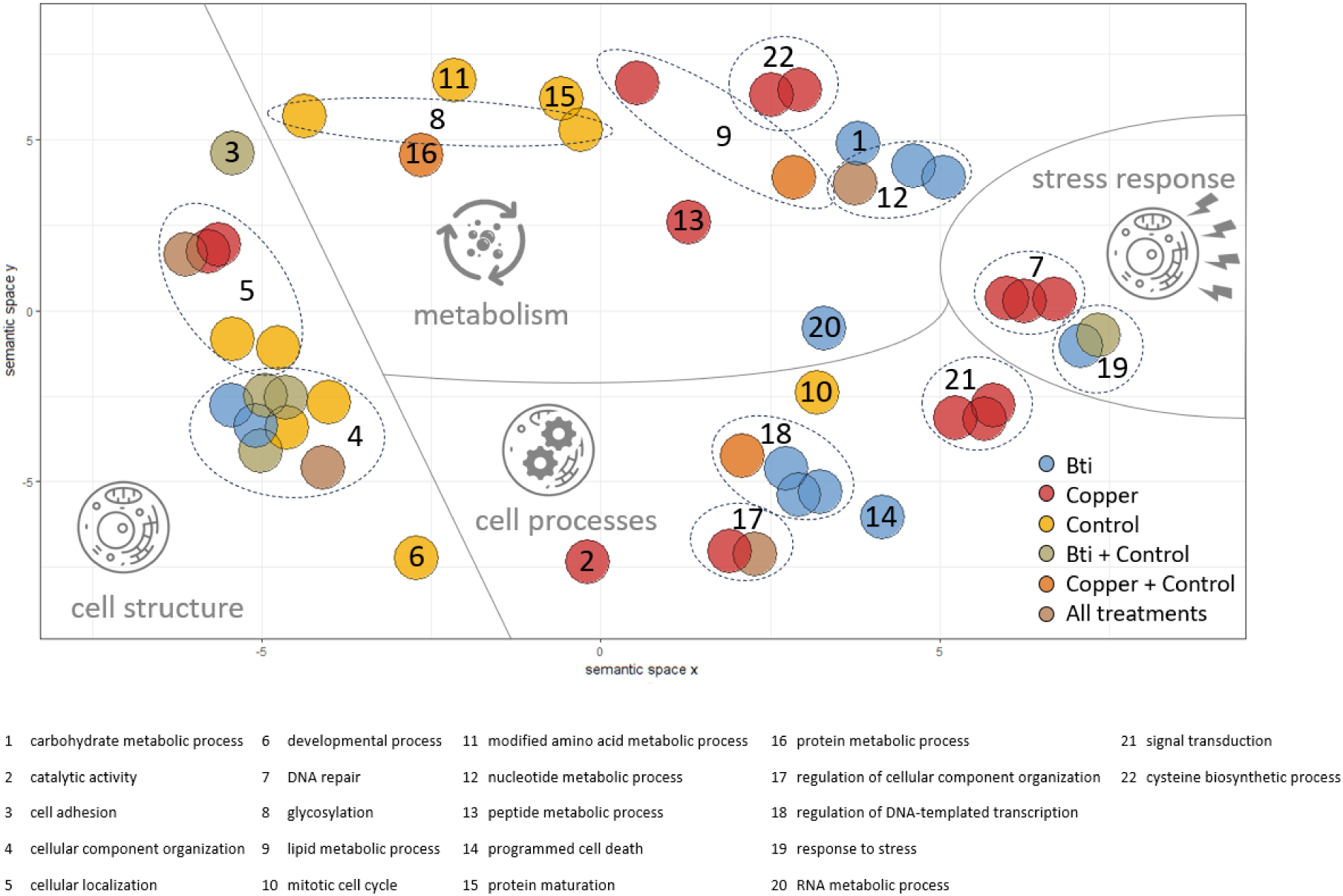
Scatterplot of 45 significantly enriched Gene Ontology (GO) terms, visualized in a two-dimensional semantic space using REVIGO (Supek et al., 2011). The x- and y-axes represent semantic space, where closer terms have higher functional similarity. Each colored circle represents a GO term, with colors indicating the treatment(s) in which the term was significantly enriched (see legend for color coding). Some functionally related GO terms are grouped within dashed-lined circles. Each colored circle or group is assigned a number, and below the figure, a corresponding list provides higher-level biological processes associated with the respective GO term(s). A full list of specific GO terms is available in the Supporting Information, Table S1.

## Discussion

In this study, we investigated the genomic response of *C. riparius* to chronic exposure to two ecologically relevant anthropogenic pollutants, with the aim of identifying early genomic signals of selection even in the absence of unambiguous phenotypic adaptation. Linkage-disequilibrium (LD) patterns indicated a bottleneck in all three treatments, with the strongest signal in the copper populations, followed by Bti, and weakest in the controls. This pattern aligns with findings by Nieto-Blázquez et al. (2025), who reported that heavy metal exposure, such as cadmium, can suppress recombination. The observed gradient in LD strength was mirrored by corresponding reductions in genome-wide nucleotide diversity (Tajima’s π), with the copper treatment exhibiting a statistically significant reduction in genome-wide nucleotide diversity. Together, these results suggest that pollutant exposure intensified demographic effects, likely due to bottlenecks, compared to the controls, creating the necessary conditions for evolutionary responses to occur.

PCA and convergence correlation analyses revealed non-parallel genome-wide allele frequency changes across treatments (Figure 1). Both convergence and divergence were observed between replicate pairs (Figure 3), regardless of treatment, indicating the absence of consistent, parallel evolution. Following Barghi et al. (2019), genetic redundancy may explain why allele frequency changes occur only in subsets of replicates, even under similar selective pressures. To account for this, we identified candidate SNPs within subsets of replicated populations (Figure 2), i.e., candidate SNPs had to be significant only in at least four out of six replicated populations. The number of unique candidate SNPs was similar across replicates, while overlap among different treatments was limited, as expected under distinct selection regimes. Interestingly, copper treatment yielded the fewest candidate SNPs despite evidence of the strongest bottleneck and a significant selection signal via covariance analysis. Importantly, there was no indication that copper and Bti treatments shared a high number of general ‘pollutant stress-response’ SNPs, supporting the conclusion of stressor-specific genomic responses. However, the number of candidate SNPs may be an overestimation, as we tested each locus individually to identify candidates. Our calculations assumed independence among loci — a simplification, given that physical linkage likely extends across several hundred base pairs. As a result, some candidate SNPs may represent linked variants within the same selective sweep rather than independent signals of selection.

Covariance-based analysis revealed that non-parallel shifts—arising from genetic drift or selection responses occurring in fewer than four replicates—explained most of the genomic variation in both the copper- and Bti-treated populations (Figure 4). Taken together with the limited evidence for consistent phenotypic adaptation (Kolbenschlag et al., 2024; Pietz et al., submitted), this suggests that selective pressures were generally weak. Nevertheless, we found clear evidence that selection contributed to genome-wide allele frequency change in the copper treatment. In contrast, no such evidence was detected in the Bti treatment, where allele frequency changes appeared to be driven primarily by neutral or replicate-specific dynamics. This apparent lack of consistent selection signals may be explained by the nature of the Bti exposure regime. Unlike copper, which was continuously present in the sediment, Bti had to be applied every two weeks due to its presumed rapid degradation and loss during medium exchange. This intermittent exposure reflects common mosquito control practices, where Bti is applied multiple times per season (Becker et al., 2018). Given that generation times in our experiment likely exceeded two weeks (Hooper et al., 2003; OECD, 2004), it is reasonable to assume that all larvae experienced Bti exposure at least once during development. However, because older larvae are known to be less sensitive to Bti than younger ones (Kästel et al., 2017), the impact on individual health likely varied with larval age at application date. This would lead to selection pressures that varied among individuals, rather than acting uniformly across the population. Nevertheless, an absence of genome-wide covariance of allele-frequency changes as seen in Bti treatments does not necessarily imply an absence of selection. Such signals can be masked by unselected sites and by opposing selection pressures at different loci (Lynch et al., 2024). Indeed, in the Bti treatment, we observed greater heterogeneity across genomic regions, as reflected in the broader confidence intervals across genomic windows. This pattern suggests that a few large-effect loci, rather than widespread polygenic adaptation, may have driven the observed changes (Supporting Information, Figure S4). This difference in the nature of the targets of selection may also explain the disagreement between number of candidate loci (Figure 2) and the covariance signals of selection for Bti and copper (Figure 4). For Bti, strong selective responses in few replicates could drive a significant Fisher’s exact test result, but would result in a weak and variable covariance signal. In contrast, a diffuse polygenic signal may be missed by the Fisher’s exact test but detected using covariance. Thus, the differences between the two results reveals the nature of the underlying genomic architecture of the response to selection for each treatment.

Although genome-wide signals of selection were heterogeneous, functional enrichment analyses revealed evolutionary responses across all treatments, supporting the conclusion that selective pressures acted in a stressor-specific manner. Stress-response and cellular processes were primarily enriched in the Bti- and copper-exposed populations, whereas GO terms related to cell structure were more prominent in controls (Figure 5). The distinct clustering of metabolically related GO terms provides further evidence that different adaptive pathways were followed in response to the specific selective environments. Notably, while Bti and copper treatments showed a similar number of enriched GO terms, there was no overlap between them, emphasizing the stressor-specific nature of these evolutionary responses to pollutant exposure.

Enrichment of the GO term “regulation of apoptotic process”, as seen in all replicates of Bti-exposed populations, suggests a role of programmed cell death in maintaining gut integrity under Bti stress, consistent with midgut restoration mechanisms observed in *Drosophila* after *Bacillus thuringiensis* exposure (Loudhaief et al., 2017). Additionally, the Bti-related enrichment of the GO term “defense response” indicates activation of immune pathways aimed at limiting damage and promoting recovery. In copper-exposed populations, enriched cysteine biosynthesis pathways highlight the well-known role of cysteine metabolism in metal detoxification (Jeppe et al., 2014, 2017). Furthermore, GO terms related to DNA damage repair are consistent with the genotoxic effects of copper exposure documented in *C. riparius* (Michailova et al., 2006). Interestingly, phototransduction pathways were also enriched, which have previously been associated with cadmium exposure in *C. riparius* (Doria et al., 2022), though the exact role of these pathways under metal stress remains speculative. In *C. riparius*, increased metal excretion and cuticle shedding have been described as responses to metal exposure and proposed as adaptive mechanisms to reduce metal burden (Postma et al., 1996; Groenendijk et al., 1999b). However, we found no enrichment of GO terms directly associated with these processes, suggesting either alternative detoxification strategies or that the observed mechanisms in natural populations reflect phenotypic plasticity rather than genetic adaptation.

In contrast, control populations showed enrichment of GO terms related to reproduction, abscission, cell cycle, and organelle functions, suggesting adaptive responses are driven by intraspecific competition and density-dependent selection in the absence of experimental stressors. This aligns with previous observations (Pfenninger & Foucault, 2020) that even under seemingly favorable laboratory conditions, selection on both life-history traits and the efficiency of basic cellular functions remains an ongoing evolutionary force.

Despite evidence for ongoing evolutionary processes at the genomic level, trait-based assessments of the evolved populations revealed limited and inconsistent phenotypic adaptation (Kolbenschlag et al., 2024; Pietz et al., submitted). A similar outcome was reported by Doria et al. (2022), who suggested that observable fitness-related phenotypic changes may require more than a few generations of selection to manifest. This idea is further supported by findings from Rigano et al. (2025), who conducted a multigeneration experiment in which *C. riparius* populations were exposed to polluted sediments and tracked across generations for both allele frequency changes and fitness traits. While shifts in allele frequencies were detectable from the first generation onward, phenotypic fitness (e.g., population growth rate) initially decreased in the first generations before showing signs of recovery— highlighting that genomic adaptation can precede detectable phenotypic responses.

Additionally, the choice of phenotypic endpoints can critically influence the detection of adaptive changes. For example, Ducrot et al. (2004) demonstrated that copper exposure in *C. riparius* reduced the proportion of emerging females capable of producing egg masses and increased egg-mass deformities. Similarly, Paris et al. (2011b) linked resistance to Bti toxins in the mosquito *Aedes aegypti* to reduced female fecundity (i.e., the number of eggs laid). Reproduction-related traits like these, reflecting actual reproductive success, were not assessed in our selection experiment. Thus, relevant adaptive phenotypes may have been overlooked. In the case of Bti, it is also possible that the adaptive processes occurring at the genetic level targeted specific toxins within the Bti mixture (cf. Paris et al., 2011a). However, the overall toxicity of the commercial Bti product, which contains multiple active components, potentially masked most measurable phenotypic effects (cf. Stalinski et al., 2014). Moreover, our trait-based assessments of the evolved populations may not have fully reflected the selective conditions during the chronic exposure, particularly since competition was substantially reduced in test vessels containing only a limited number of organisms. Finally, biological variability likely limited the detection of subtle phenotypic changes; increased replication and more sensitive assays (e.g., transcriptomic profiling or enzymatic activity assays) may be needed to reveal early-stage phenotypic adaptations.

## Conclusion

Taken together, our results suggest that, while non-parallel shifts dominated genome-wide allele frequency dynamics, selection has started to modestly shape treatment-specific adaptive responses. The observed patterns imply that adaptation to chronic pollutant exposure in *C. riparius* may initially proceed through subtle, pathway-specific modifications rather than through large-scale, parallel genomic shifts. Our study highlights the complexity of predicting adaptive responses to environmental pollutants. Secondary Evolve and Resequencing studies, where evolved populations are crossed back with the ancestral population and re-exposed to the same selective environment, could validate weak or replicate-specific selection signals (Burny et al., 2020). Additionally, computational simulations can aid in optimizing experimental designs, for instance by identifying the appropriate number of replicates, generations of selection, and starting haplotype diversity to robustly capture adaptive genomic responses (Vlachos & Kofler, 2018). Furthermore, integrating transcriptomic data (Doria et al., 2022) could provide functional insights by revealing gene expression changes associated with adaptive responses. Such complementary approaches may confirm candidate genes or uncover adaptive mechanisms that SNP-based analyses alone might overlook.

Understanding evolutionary responses is essential for predicting long-term ecological consequences, particularly in the face of persistent environmental pollution. Traditional ecotoxicological assessments predominantly focus on short-term toxicity (Padilla Suarez et al., 2023; Thoré et al., 2021) and acute effects on survival (Straub et al., 2020), often overlooking the potential for adaptive responses over multiple generations. Evolve and Resequencing approaches bridge this gap by integrating experimental evolution with genomics, providing direct insights into adaptive genetic change. This enables the detection of subtle or early-stage adaptations that may not yet manifest at the phenotypic level. In this way, by assessing population-level evolutionary responses to diverse pollutants, such as biocides and heavy metals, Evolve and Resequencing studies offer valuable empirical data on both the capacity and constraints of adaptation. However, translating laboratory-based findings to natural populations remains challenging (Phillips & Burke, 2021). Differences in generation times, effective population sizes, standing genetic variation, and environmental conditions — all of which can profoundly shape evolutionary trajectories — must be considered. Comparisons with natural populations are therefore essential to contextualize laboratory observations. However, despite its laboratory-based nature, this study lays important groundwork for elucidating evolutionary processes driven by pollution in riparian systems.

## Data availability

The sequence data generated in this study are available from the National Center for Biotechnology Information (NCBI) under the BioProject accession number PRJNA1282899. Code to run all analyses can be found on Zenodo: https://doi.org/10.5281/zenodo.15744395. Construction protocols for the chironomid-handling tools developed in this study are available on protocols.io: https://dx.doi.org/10.17504/protocols.io.3byl49qorgo5/v1.

## Acknowledgements

We thank Verena Gerstle and Lukas Beyer for their assistance with larval counts, and Anja Knäbel for her continuous support throughout the study. We are grateful to Barbara Feldmeyer for her valuable guidance on sequencing and data analysis. We also thank ECT, BASF and the Molecular Ecology team at the Senckenberg Biodiversity and Climate Research Centre for providing chironomid egg masses. This study was funded by the Deutsche Forschungsgemeinschaft (DFG, German Research Foundation) – Research Training Group SystemLink 326210499/GRK2360.

## Author Contributions

NR, SK, SP, MB, and KS conceived and designed the study. KS, MB, and MP secured funding for the project. NR, SK, and SP conducted the experiments and collected the samples. NR processed the samples. Visualizations and code were based on previous work by RB. NR performed the bioinformatic and statistical analyses with support from RB and MP. NR wrote the first draft of the manuscript. All authors contributed to manuscript revisions and approved the final version.

## Ethics Statement

This study did not involve work with vertebrate animals, endangered species, or human participants, and thus did not require formal ethical approval. All experimental procedures complied with local regulations and institutional guidelines.

