## Supporting Information for "Pollution-Driven Selection in Riparian Ecosystems: Genome-Wide Responses to *Bacillus thuringiensis israelensis* and Copper in a Non-biting Midge"

#### List of contents

#### Figure S1

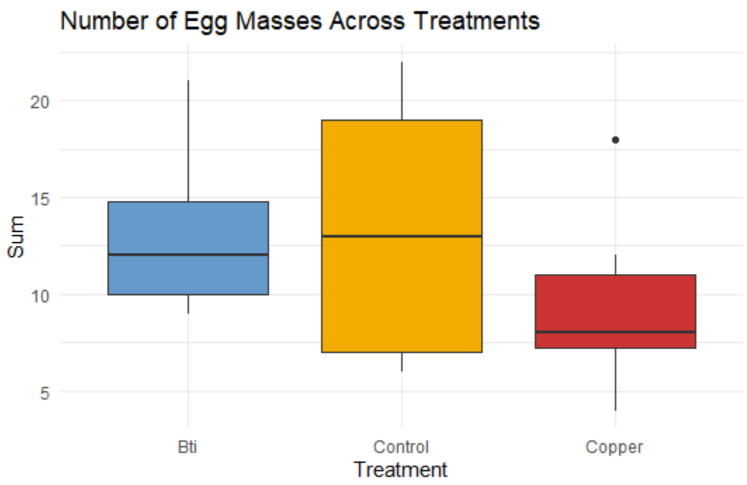

*Figure S1 Number of egg masses sampled per treatment (n = 6) following 26 weeks of chronic exposure. Egg masses were collected over three consecutive days and stored in SAM-5S medium until hatching. Larvae hatching from these egg masses were used for population genomic analyses. Boxplots indicate the interquartile range (IQR), horizontal lines denote the median, whiskers extend to 1.5× IQR, and points beyond this range are shown as outliers.*

#### Figure S2

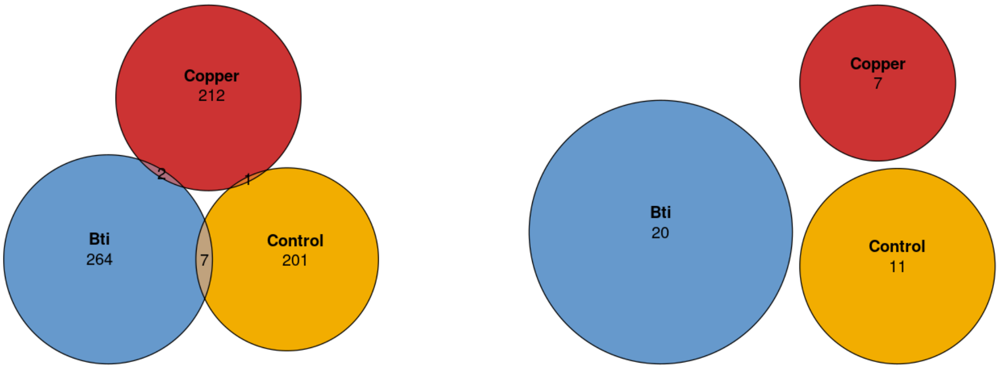

*Figure S2 Venn diagrams showing the number of candidate SNPs under stricter significance thresholds: (left) ≥5 of 6 replicates, and (right) all 6 replicates per treatment. Stricter criteria greatly reduced both treatment-specific and shared SNPs.*

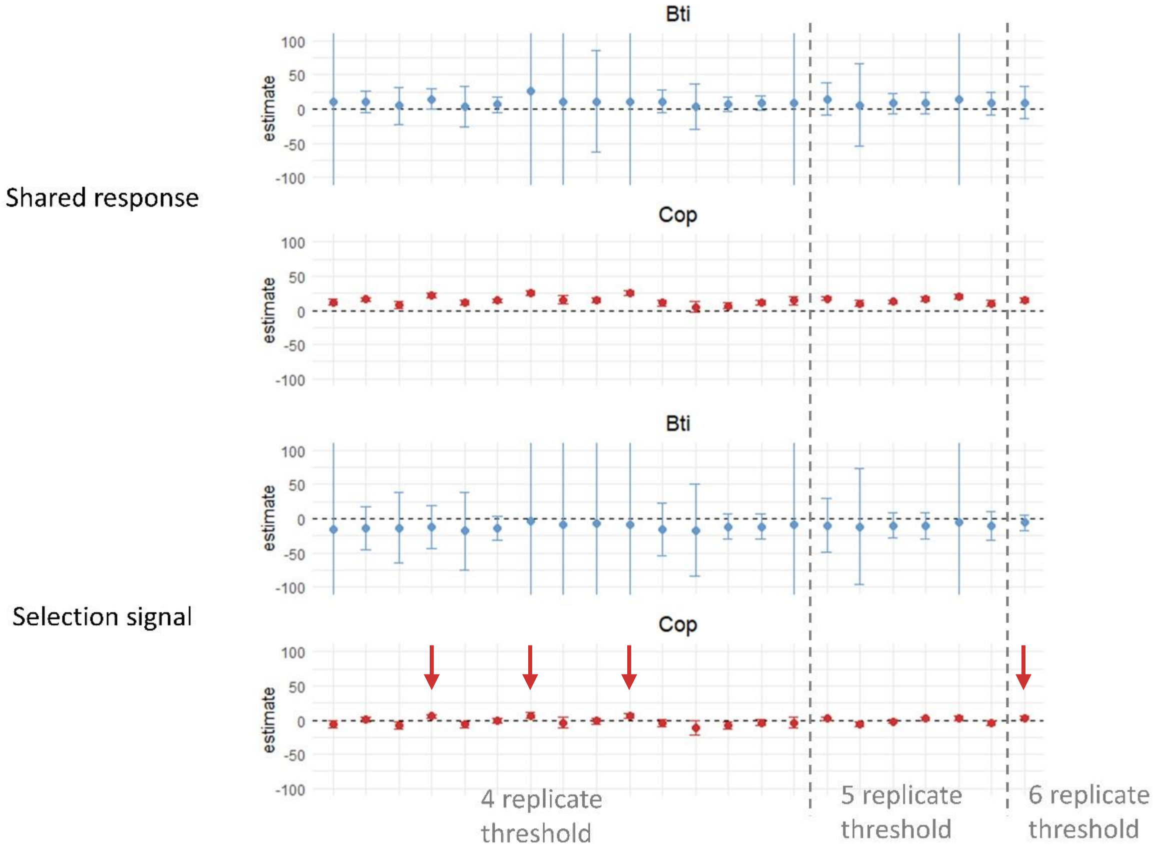

*Figure S3 Partitioning of allele frequency change variance into shared and selection components across replicate combinations for Bti- and copper-treated populations. Upper panels show the percentage of total variance attributed to shared allele frequency change (i.e., parallel change), and lower panels show the variance attributed to selection after accounting for laboratory adaptation. Statistically significant positive selection estimates are highlighted with arrows. Each point represents a replicate combination (15 four-replicate, 6 five-replicate, and 1 six-replicate combination per treatment), with 95% confidence intervals indicated by error bars. Variance components are expressed as percentages of total variance in allele frequency change from F0 to F8. Error bars extending beyond the plotting range (±100%) are truncated visually.*

### Figure S4

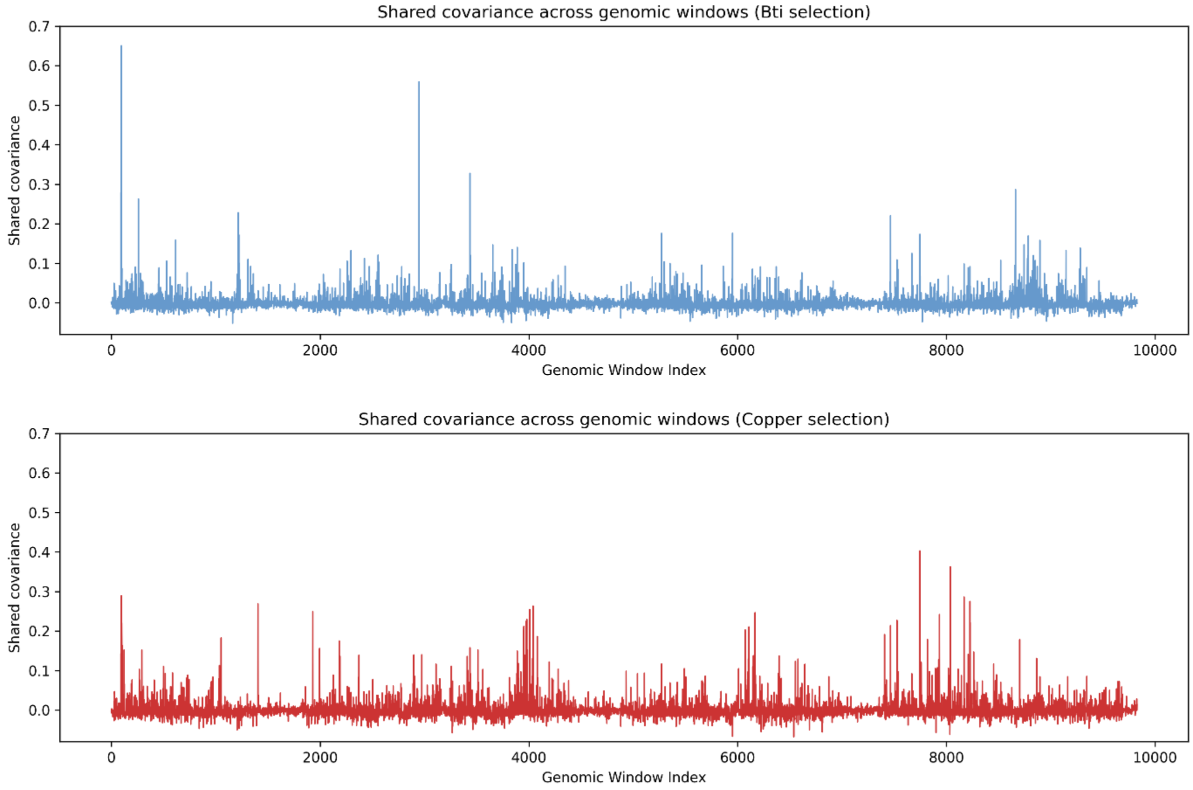

*Figure S4 Genome-wide distribution of shared covariance across genomic windows in six-out-of-six* Chironomus riparius *populations exposed to Bti (top) and copper (bottom). For each genomic window, the proportion of total genetic variance attributable to shared allele frequency changes across replicates is shown.*

### Table S1

*Table S1 Summary statistics of variance partitioning results for all tested replicate combinations across treatments. Each row reports the estimate and confidence interval (lower and upper error bounds) for one of four variance components—total variance in allele frequency change, the shared response, the portion attributable to laboratory adaptation, and the inferred contribution of experimental selection—for a specific replicate combination in either Bti-treated or copper-exposed populations. The combination number refers to a specific set of replicates, and the number of replicates used in each is indicated. The range column reflects the span between the upper and lower error bounds. See Supplementary Figure S2 for graphical visualization of the shared and selection variance components.*

| Sort | treatment | variable | combination number | lower_err | estimate | upper_err | range | based on .. replicates |
|---|---|---|---|---|---|---|---|---|
| 1 | Bti | total | 1 | 0.0169 | 0.0178 | 0.0188 | 0.0019 | 4 |
| 2 | Bti | shared | 1 | -618.215 | 10.951 | 640.118 | 1258.333 | 4 |
| 3 | Bti | lab | 1 | -1147.781 | 26.757 | 1201.296 | 2349.077 | 4 |
| 4 | Bti | selection | 1 | -1818.411 | -15.806 | 1786.799 | 3605.21 | 4 |
| 5 | Bti | total | 2 | 0.0198 | 0.0209 | 0.022 | 0.0022 | 4 |
| 6 | Bti | shared | 2 | -4.19 | 11.295 | 26.78 | 30.97 | 4 |
| 7 | Bti | lab | 2 | 4.008 | 24.414 | 44.82 | 40.812 | 4 |
| 8 | Bti | selection | 2 | -44.733 | -13.119 | 18.496 | 63.229 | 4 |
| 9 | Bti | total | 3 | 0.0196 | 0.0206 | 0.0216 | 0.002 | 4 |
| 10 | Bti | shared | 3 | -21.682 | 4.925 | 31.531 | 53.213 | 4 |
| 11 | Bti | lab | 3 | -10.192 | 18.075 | 46.341 | 56.533 | 4 |
| 12 | Bti | selection | 3 | -64.265 | -13.15 | 37.965 | 102.23 | 4 |
| 13 | Bti | total | 4 | 0.0196 | 0.0207 | 0.0217 | 0.0021 | 4 |

| | | | | | | | | |
|---|---|---|---|---|---|---|---|---|
| 14 | Bti | shared | 4 | 0.519 | 15.037 | 29.555 | 29.036 | 4 |
| 15 | Bti | lab | 4 | 4.995 | 27.112 | 49.229 | 44.234 | 4 |
| 16 | Bti | selection | 4 | -44.032 | -12.074 | 19.883 | 63.915 | 4 |
| 17 | Bti | total | 5 | 0.0194 | 0.0204 | 0.0214 | 0.002 | 4 |
| 18 | Bti | shared | 5 | -26.713 | 3.348 | 33.409 | 60.122 | 4 |
| 19 | Bti | lab | 5 | -9.36 | 20.738 | 50.836 | 60.196 | 4 |
| 20 | Bti | selection | 5 | -73.975 | -17.39 | 39.195 | 113.17 | 4 |
| 21 | Bti | total | 6 | 0.0224 | 0.0235 | 0.0246 | 0.0022 | 4 |
| 22 | Bti | shared | 6 | -5.543 | 6.489 | 18.521 | 24.064 | 4 |
| 23 | Bti | lab | 6 | 9.459 | 19.443 | 29.426 | 19.967 | 4 |
| 24 | Bti | selection | 6 | -30.425 | -12.953 | 4.518 | 34.943 | 4 |
| 25 | Bti | total | 7 | 0.014 | 0.015 | 0.0159 | 0.0019 | 4 |
| 26 | Bti | shared | 7 | -6455.978 | 26.603 | 6509.184 | 12965.162 | 4 |
| 27 | Bti | lab | 7 | -206971.779 | 29.687 | 207031.152 | 414002.931 | 4 |
| 28 | Bti | selection | 7 | -203467.4 | -3.084 | 203461.232 | 406928.632 | 4 |
| 29 | Bti | total | 8 | 0.0138 | 0.0147 | 0.0155 | 0.0017 | 4 |
| 30 | Bti | shared | 8 | -4093.184 | 10.619 | 4114.422 | 8207.606 | 4 |
| 31 | Bti | lab | 8 | -12646.692 | 18.289 | 12683.27 | 25329.962 | 4 |
| 32 | Bti | selection | 8 | -16774.891 | -7.671 | 16759.55 | 33534.441 | 4 |
| 33 | Bti | total | 9 | 0.0168 | 0.0178 | 0.0187 | 0.0019 | 4 |
| 34 | Bti | shared | 9 | -63.186 | 11.572 | 86.33 | 149.516 | 4 |
| 35 | Bti | lab | 9 | -163.396 | 18.1 | 199.597 | 362.993 | 4 |
| 36 | Bti | selection | 9 | -258.171 | -6.528 | 245.115 | 503.286 | 4 |
| 37 | Bti | total | 10 | 0.0166 | 0.0175 | 0.0185 | 0.0019 | 4 |
| 38 | Bti | shared | 10 | -150.599 | 10.841 | 172.281 | 322.88 | 4 |
| 39 | Bti | lab | 10 | -283.03 | 19.377 | 321.785 | 604.815 | 4 |
| 40 | Bti | selection | 10 | -469.503 | -8.537 | 452.43 | 921.933 | 4 |
| 41 | Bti | total | 11 | 0.019 | 0.02 | 0.021 | 0.002 | 4 |
| 42 | Bti | shared | 11 | -4.597 | 11.452 | 27.5 | 32.097 | 4 |
| 43 | Bti | lab | 11 | 0.442 | 26.251 | 52.059 | 51.617 | 4 |
| 44 | Bti | selection | 11 | -53.086 | -14.799 | 23.488 | 76.574 | 4 |
| 45 | Bti | total | 12 | 0.0187 | 0.0197 | 0.0207 | 0.002 | 4 |
| 46 | Bti | shared | 12 | -29.799 | 3.26 | 36.32 | 66.119 | 4 |
| 47 | Bti | lab | 12 | -18.091 | 19.644 | 57.379 | 75.47 | 4 |
| 48 | Bti | selection | 12 | -84.125 | -16.384 | 51.357 | 135.482 | 4 |
| 49 | Bti | total | 13 | 0.0217 | 0.0228 | 0.0239 | 0.0022 | 4 |
| 50 | Bti | shared | 13 | -3.959 | 7.181 | 18.321 | 22.28 | 4 |
| 51 | Bti | lab | 13 | 7.722 | 18.458 | 29.194 | 21.472 | 4 |
| 52 | Bti | selection | 13 | -29.486 | -11.277 | 6.933 | 36.419 | 4 |
| 53 | Bti | total | 14 | 0.0215 | 0.0226 | 0.0236 | 0.0021 | 4 |
| 54 | Bti | shared | 14 | -0.998 | 9.581 | 20.16 | 21.158 | 4 |
| 55 | Bti | lab | 14 | 9.454 | 20.868 | 32.283 | 22.829 | 4 |
| 56 | Bti | selection | 14 | -29.321 | -11.288 | 6.745 | 36.066 | 4 |
| 57 | Bti | total | 15 | 0.0159 | 0.0169 | 0.0178 | 0.0019 | 4 |
| 58 | Bti | shared | 15 | -502.029 | 9.807 | 521.643 | 1023.672 | 4 |

| | | | | | | | | |
|---|---|---|---|---|---|---|---|---|
| 59 | Bti | lab | 15 | -1207.532 | 18.043 | 1243.618 | 2451.15 | 4 |
| 60 | Bti | selection | 15 | -1741.909 | -8.236 | 1725.438 | 3467.347 | 4 |
| 61 | Cop | total | 1 | 0.0275 | 0.0288 | 0.0301 | 0.0026 | 4 |
| 62 | Cop | shared | 1 | 7.306 | 11.841 | 16.377 | 9.071 | 4 |
| 63 | Cop | lab | 1 | 13.779 | 17.589 | 21.398 | 7.619 | 4 |
| 64 | Cop | selection | 1 | -10.937 | -5.747 | -0.557 | 10.38 | 4 |
| 65 | Cop | total | 2 | 0.032 | 0.0333 | 0.0347 | 0.0027 | 4 |
| 66 | Cop | shared | 2 | 12.548 | 15.866 | 19.184 | 6.636 | 4 |
| 67 | Cop | lab | 2 | 12.577 | 15.375 | 18.173 | 5.596 | 4 |
| 68 | Cop | selection | 2 | -2.73 | 0.492 | 3.713 | 6.443 | 4 |
| 69 | Cop | total | 3 | 0.0261 | 0.0273 | 0.0285 | 0.0024 | 4 |
| 70 | Cop | shared | 3 | 2.629 | 7.509 | 12.388 | 9.759 | 4 |
| 71 | Cop | lab | 3 | 12.339 | 16.06 | 19.781 | 7.442 | 4 |
| 72 | Cop | selection | 3 | -13.67 | -8.551 | -3.432 | 10.238 | 4 |
| 73 | Cop | total | 4 | 0.0339 | 0.0353 | 0.0367 | 0.0028 | 4 |
| 74 | Cop | shared | 4 | 18.714 | 21.813 | 24.913 | 6.199 | 4 |
| 75 | Cop | lab | 4 | 14.077 | 16.484 | 18.892 | 4.815 | 4 |
| 76 | Cop | selection | 4 | 2.214 | 5.329 | 8.444 | 6.23 | 4 |
| 77 | Cop | total | 5 | 0.0281 | 0.0293 | 0.0305 | 0.0024 | 4 |
| 78 | Cop | shared | 5 | 7.036 | 11.032 | 15.028 | 7.992 | 4 |
| 79 | Cop | lab | 5 | 14.15 | 17.245 | 20.34 | 6.19 | 4 |
| 80 | Cop | selection | 5 | -10.655 | -6.213 | -1.772 | 8.883 | 4 |
| 81 | Cop | total | 6 | 0.0325 | 0.0338 | 0.0352 | 0.0027 | 4 |
| 82 | Cop | shared | 6 | 11.077 | 14.155 | 17.232 | 6.155 | 4 |
| 83 | Cop | lab | 6 | 12.678 | 14.953 | 17.228 | 4.55 | 4 |
| 84 | Cop | selection | 6 | -3.749 | -0.798 | 2.153 | 5.902 | 4 |
| 85 | Cop | total | 7 | 0.0299 | 0.0312 | 0.0325 | 0.0026 | 4 |
| 86 | Cop | shared | 7 | 21.421 | 25.238 | 29.055 | 7.634 | 4 |
| 87 | Cop | lab | 7 | 15.456 | 18.931 | 22.405 | 6.949 | 4 |
| 88 | Cop | selection | 7 | 1.969 | 6.307 | 10.645 | 8.676 | 4 |
| 89 | Cop | total | 8 | 0.0241 | 0.0252 | 0.0264 | 0.0023 | 4 |
| 90 | Cop | shared | 8 | 10.192 | 15.658 | 21.123 | 10.931 | 4 |
| 91 | Cop | lab | 8 | 14.168 | 19.357 | 24.546 | 10.378 | 4 |
| 92 | Cop | selection | 8 | -11.233 | -3.699 | 3.835 | 15.068 | 4 |
| 93 | Cop | total | 9 | 0.0285 | 0.0297 | 0.031 | 0.0025 | 4 |
| 94 | Cop | shared | 9 | 11.639 | 15.39 | 19.142 | 7.503 | 4 |
| 95 | Cop | lab | 9 | 13.417 | 16.608 | 19.799 | 6.382 | 4 |
| 96 | Cop | selection | 9 | -5.41 | -1.217 | 2.975 | 8.385 | 4 |
| 97 | Cop | total | 10 | 0.0304 | 0.0317 | 0.033 | 0.0026 | 4 |
| 98 | Cop | shared | 10 | 21.447 | 24.86 | 28.274 | 6.827 | 4 |
| 99 | Cop | lab | 10 | 15.332 | 18.15 | 20.968 | 5.636 | 4 |
| 100 | Cop | selection | 10 | 3.133 | 6.711 | 10.288 | 7.155 | 4 |
| 101 | Cop | total | 11 | 0.0281 | 0.0293 | 0.0306 | 0.0025 | 4 |
| 102 | Cop | shared | 11 | 6.433 | 10.954 | 15.475 | 9.042 | 4 |
| 103 | Cop | lab | 11 | 12.436 | 16.116 | 19.796 | 7.36 | 4 |

| | | | | | | | | |
|---|---|---|---|---|---|---|---|---|
| 104 | Cop | selection | 11 | -10.335 | -5.162 | 0.011 | 10.346 | 4 |
| 105 | Cop | total | 12 | 0.0223 | 0.0234 | 0.0244 | 0.0021 | 4 |
| 106 | Cop | shared | 12 | -3.099 | 4.864 | 12.827 | 15.926 | 4 |
| 107 | Cop | lab | 12 | 9.757 | 15.854 | 21.95 | 12.193 | 4 |
| 108 | Cop | selection | 12 | -21.689 | -10.99 | -0.291 | 21.398 | 4 |
| 109 | Cop | total | 13 | 0.0267 | 0.0279 | 0.0291 | 0.0024 | 4 |
| 110 | Cop | shared | 13 | 2.21 | 6.801 | 11.392 | 9.182 | 4 |
| 111 | Cop | lab | 13 | 10.492 | 13.943 | 17.395 | 6.903 | 4 |
| 112 | Cop | selection | 13 | -12.264 | -7.142 | -2.021 | 10.243 | 4 |
| 113 | Cop | total | 14 | 0.0286 | 0.0299 | 0.0311 | 0.0025 | 4 |
| 114 | Cop | shared | 14 | 7.247 | 11.285 | 15.323 | 8.076 | 4 |
| 115 | Cop | lab | 14 | 12.608 | 15.539 | 18.471 | 5.863 | 4 |
| 116 | Cop | selection | 14 | -8.636 | -4.254 | 0.128 | 8.764 | 4 |
| 117 | Cop | total | 15 | 0.0246 | 0.0258 | 0.0269 | 0.0023 | 4 |
| 118 | Cop | shared | 15 | 8.409 | 14.136 | 19.862 | 11.453 | 4 |
| 119 | Cop | lab | 15 | 13.106 | 18.133 | 23.16 | 10.054 | 4 |
| 120 | Cop | selection | 15 | -11.576 | -3.997 | 3.582 | 15.158 | 4 |
| 121 | Bti | total | 1 | 0.0179 | 0.0189 | 0.0198 | 0.0019 | 5 |
| 122 | Bti | shared | 1 | -8.773 | 14.51 | 37.794 | 46.567 | 5 |
| 123 | Bti | lab | 1 | 3.593 | 24.236 | 44.879 | 41.286 | 5 |
| 124 | Bti | selection | 1 | -48.684 | -9.726 | 29.233 | 77.917 | 5 |
| 125 | Bti | total | 2 | 0.0177 | 0.0186 | 0.0196 | 0.0019 | 5 |
| 126 | Bti | shared | 2 | -53.939 | 6.276 | 66.491 | 120.43 | 5 |
| 127 | Bti | lab | 2 | -11.454 | 17.763 | 46.98 | 58.434 | 5 |
| 128 | Bti | selection | 2 | -95.967 | -11.487 | 72.993 | 168.96 | 5 |
| 129 | Bti | total | 3 | 0.0201 | 0.0211 | 0.0221 | 0.002 | 5 |
| 130 | Bti | shared | 3 | -7.507 | 8.14 | 23.786 | 31.293 | 5 |
| 131 | Bti | lab | 3 | 9.2 | 17.686 | 26.171 | 16.971 | 5 |
| 132 | Bti | selection | 3 | -28.138 | -9.546 | 9.047 | 37.185 | 5 |
| 133 | Bti | total | 4 | 0.02 | 0.0209 | 0.0219 | 0.0019 | 5 |
| 134 | Bti | shared | 4 | -7.027 | 8.961 | 24.95 | 31.977 | 5 |
| 135 | Bti | lab | 4 | 10.241 | 18.768 | 27.295 | 17.054 | 5 |
| 136 | Bti | selection | 4 | -28.699 | -9.807 | 9.084 | 37.783 | 5 |
| 137 | Bti | total | 5 | 0.0155 | 0.0164 | 0.0172 | 0.0017 | 5 |
| 138 | Bti | shared | 5 | -15437.727 | 13.629 | 15464.984 | 30902.711 | 5 |
| 139 | Bti | lab | 5 | -17149.623 | 19.281 | 17188.185 | 34337.808 | 5 |
| 140 | Bti | selection | 5 | -32608.194 | -5.652 | 32596.889 | 65205.083 | 5 |
| 141 | Bti | total | 6 | 0.0194 | 0.0204 | 0.0214 | 0.002 | 5 |
| 142 | Bti | shared | 6 | -8.315 | 8.226 | 24.767 | 33.082 | 5 |
| 143 | Bti | lab | 6 | 8.889 | 17.913 | 26.937 | 18.048 | 5 |
| 144 | Bti | selection | 6 | -30.387 | -9.687 | 11.012 | 41.399 | 5 |
| 145 | Cop | total | 1 | 0.0303 | 0.0316 | 0.0329 | 0.0026 | 5 |
| 146 | Cop | shared | 1 | 14.18 | 17.401 | 20.621 | 6.441 | 5 |
| 147 | Cop | lab | 1 | 12.669 | 15.178 | 17.688 | 5.019 | 5 |
| 148 | Cop | selection | 1 | -0.637 | 2.222 | 5.081 | 5.718 | 5 |

| | | | | | | | | |
|---|---|---|---|---|---|---|---|---|
| 149 | Cop | total | 2 | 0.0257 | 0.0268 | 0.028 | 0.0023 | 5 |
| 150 | Cop | shared | 2 | 5.851 | 10.283 | 14.715 | 8.864 | 5 |
| 151 | Cop | lab | 2 | 12.692 | 15.836 | 18.98 | 6.288 | 5 |
| 152 | Cop | selection | 2 | -9.348 | -5.553 | -1.758 | 7.59 | 5 |
| 153 | Cop | total | 3 | 0.0292 | 0.0304 | 0.0316 | 0.0024 | 5 |
| 154 | Cop | shared | 3 | 9.01 | 12.229 | 15.448 | 6.438 | 5 |
| 155 | Cop | lab | 3 | 11.647 | 14.136 | 16.625 | 4.978 | 5 |
| 156 | Cop | selection | 3 | -4.534 | -1.907 | 0.721 | 5.255 | 5 |
| 157 | Cop | total | 4 | 0.0308 | 0.032 | 0.0332 | 0.0024 | 5 |
| 158 | Cop | shared | 4 | 13.873 | 16.859 | 19.845 | 5.972 | 5 |
| 159 | Cop | lab | 4 | 12.608 | 14.72 | 16.832 | 4.224 | 5 |
| 160 | Cop | selection | 4 | -0.393 | 2.139 | 4.671 | 5.064 | 5 |
| 161 | Cop | total | 5 | 0.0276 | 0.0287 | 0.0299 | 0.0023 | 5 |
| 162 | Cop | shared | 5 | 15.823 | 19.443 | 23.063 | 7.24 | 5 |
| 163 | Cop | lab | 5 | 13.728 | 16.48 | 19.232 | 5.504 | 5 |
| 164 | Cop | selection | 5 | -0.383 | 2.963 | 6.309 | 6.692 | 5 |
| 165 | Cop | total | 6 | 0.0261 | 0.0272 | 0.0284 | 0.0023 | 5 |
| 166 | Cop | shared | 6 | 5.388 | 9.734 | 14.08 | 8.692 | 5 |
| 167 | Cop | lab | 6 | 11.53 | 14.594 | 17.658 | 6.128 | 5 |
| 168 | Cop | selection | 6 | -8.62 | -4.86 | -1.1 | 7.52 | 5 |
| 169 | Bti | total | 1 | 0.0185 | 0.0194 | 0.0203 | 0.0018 | 6 |
| 170 | Bti | shared | 1 | -14.464 | 9.809 | 34.082 | 48.546 | 6 |
| 171 | Bti | lab | 1 | -1.771 | 15.415 | 32.6 | 34.371 | 6 |
| 172 | Bti | selection | 1 | -16.954 | -5.606 | 5.743 | 22.697 | 6 |
| 173 | Cop | total | 1 | 0.0283 | 0.0295 | 0.0306 | 0.0023 | 6 |
| 174 | Cop | shared | 1 | 11.283 | 14.484 | 17.686 | 6.403 | 6 |
| 175 | Cop | lab | 1 | 8.186 | 11.153 | 14.119 | 5.933 | 6 |
| 176 | Cop | selection | 1 | 0.835 | 3.332 | 5.829 | 4.994 | 6 |

#### Table S2

*Table S2 Results of Gene Ontology (GO) enrichment analysis for loci showing significant allele frequency changes after chronic exposure across treatments. The table lists the 45 enriched biological process GO terms identified in Bti-treated, copper-exposed, and control populations, including those shared across treatments. Terms were detected across replicate combinations showing statistically significant shared responses, with the corresponding number of replicates indicated.*

| Sort | GO ID | GO term | treatment | based on … replicates |
|---|---|---|---|---|
| 1 | GO:0016358 | dendrite development | all | 6 |
| 2 | GO:0031114 | regulation of microtubule depolymerization | all | 5 |
| 3 | GO:0007097 | nuclear migration | all | 4 |
| 4 | GO:0006196 | AMP catabolic process | all | 4 |
| 5 | GO:0042981 | regulation of apoptotic process | Bti | 6 |
| 6 | GO:0006952 | defense response | Bti | 5 |
| 7 | GO:0000122 | negative regulation of transcription by RNA polymerase II | Bti | 5 |
| 8 | GO:0045893 | positive regulation of DNA-templated transcription | Bti | 4 |
| 9 | GO:0006338 | chromatin remodeling | Bti | 4 |
| 10 | GO:0006741 | NADP biosynthetic process | Bti | 4 |

| | | | | |
|---|---|---|---|---|
| 11 | GO:0019674 | NAD metabolic process | Bti | 4 |
| 12 | GO:0006997 | nucleus organization | Bti | 4 |
| 13 | GO:0006357 | regulation of transcription by RNA polymerase II | Bti | 4 |
| 14 | GO:0033499 | galactose catabolic process via UDP-galactose | Bti | 4 |
| 15 | GO:0000184 | nuclear-transcribed mRNA catabolic process, nonsense-mediated decay | Bti | 4 |
| 16 | GO:0007602 | phototransduction | Copper | 5 |
| 17 | GO:0046513 | ceramide biosynthetic process | Copper | 5 |
| 18 | GO:0006606 | protein import into nucleus | Copper | 5 |
| 19 | GO:0043171 | peptide catabolic process | Copper | 5 |
| 20 | GO:0000012 | single strand break repair | Copper | 4 |
| 21 | GO:0006535 | cysteine biosynthetic process from serine | Copper | 4 |
| 22 | GO:1902275 | regulation of chromatin organization | Copper | 4 |
| 23 | GO:0008543 | fibroblast growth factor receptor signaling pathway | Copper | 4 |
| 24 | GO:0007190 | activation of adenylate cyclase activity | Copper | 4 |
| 25 | GO:0019343 | cysteine biosynthetic process via cystathionine | Copper | 4 |
| 26 | GO:0006303 | double-strand break repair via nonhomologous end joining | Copper | 4 |
| 27 | GO:0061512 | protein localization to cilium | Copper | 4 |
| 28 | GO:0051295 | establishment of meiotic spindle localization | Control | 5 |
| 29 | GO:0061077 | chaperone-mediated protein folding | Control | 5 |
| 30 | GO:0006487 | protein N-linked glycosylation | Control | 5 |
| 31 | GO:0007051 | spindle organization | Control | 5 |
| 32 | GO:0044878 | mitotic cytokinesis checkpoint signaling | Control | 4 |
| 33 | GO:0046901 | tetrahydrofolylpolyglutamate biosynthetic process | Control | 4 |
| 34 | GO:0009838 | abscission | Control | 4 |
| 35 | GO:0032979 | protein insertion into mitochondrial inner membrane from matrix | Control | 4 |
| 36 | GO:0070085 | glycosylation | Control | 4 |
| 37 | GO:0006325 | chromatin organization | Control | 4 |
| 38 | GO:0007156 | homophilic cell adhesion via plasma membrane adhesion molecules | Control-Bti | 5 |
| 39 | GO:0045104 | intermediate filament cytoskeleton organization | Control-Bti | 4 |
| 40 | GO:0042060 | wound healing | Control-Bti | 4 |
| 41 | GO:0031122 | cytoplasmic microtubule organization | Control-Bti | 4 |
| 42 | GO:0000492 | box C/D snoRNP assembly | Control-Bti | 4 |
| 43 | GO:0034244 | negative regulation of transcription elongation by RNA polymerase II | Control-Copper | 5 |
| 44 | GO:0006685 | sphingomyelin catabolic process | Control-Copper | 5 |
| 45 | GO:0019919 | peptidyl-arginine methylation, to asymmetrical-dimethyl arginine | Control-Copper | 4 |
